# N-Cadherin/α-Catenin Drive Adhesion and Actin Regulation to Orchestrate Tunneling Nanotube Formation

**DOI:** 10.1101/2025.06.27.661900

**Authors:** Sevan Belian, Anna Pepe, Roberto Notario Manzano, Anna Sartori-Rupp, Christel Brou, Chiara Zurzolo

## Abstract

Cell-to-cell communication is essential for maintaining homeostasis in multicellular organisms. Tunneling nanotubes (TNTs)—actin-based membranous connections—mediate the exchange of diverse cargoes between distant cells. Unlike other cellular protrusions, TNTs exhibit unique ultrastructural features and are enriched in the adhesion molecule N-Cadherin.

Here, we dissect the role of N-Cadherin in the formation and function of TNTs in SH-SY5Y human neuronal-like cells. We show that N-Cadherin, via its effectors α-Catenin and p120-Catenin, is a central regulator of TNT architecture and their cargo transfer capability. Regulators of cortical tension p120-Catenin, ROCK, and non-muscle myosin II also emerge as critical for TNT functionality, highlighting a mechanosensitive component to TNT regulation. Moreover, we reveal that NMIIA can be processive inside TNTs and transfer through them using actin’s retrograde flow. Finally, we identify the Cdc42–IRSp53–N-WASP pathway as a downstream effector axis enhancing intercellular transfer downstream of N-Cadherin.

Together, our findings uncover a structural and functional link between N-Cadherin signaling and TNT-mediated intercellular communication.

## Introduction

Tunneling nanotubes (TNTs) are non-adherent F-actin based membranous structures that form continuous cytoplasmic bridges between cells, spanning distances from hundreds of nanometers to up to 100 µm^1^. These intercellular connections are implicated in both physiological and pathological contexts^2–6^, facilitating the transfer of various cargoes^6–10^, including organelles, pathogens^11–14^, and misfolded proteins^7,15–18^. The defining feature of TNTs is their ability to mediate the direct transfer of a broad range of cargoes through an open cytoplasmic channel between two cells, distinguishing them from other types of cellular protrusions. Despite extensive evidence supporting their role in intercellular communication, the mechanisms underlying TNT formation, as well as the molecular components and regulatory factors involved in their structure, remain poorly understood^1^. Recently, by developing a correlative light and cryo-electron microscopy (cryo-CLEM) and tomography workflow in murine and human neuronal cell lines, we demonstrated that TNTs are structurally different from filopodia^19^. While they typically appear as single connections under fluorescence microscopy (FM), nanoscopic analysis revealed that most of them are comprised of a bundle of individual Tunneling Nanotubes (iTNTs) that can contain vesicles and organelles and can be open-ended. This data also revealed long threads coiling around the iTNT bundles as in the process of holding them together.

Notably, using immunogold labeling in mouse neuronal cells, we found the transmembrane protein N-Cadherin localized at the attachment point of these threads on the iTNTs membrane, as well as on short connections (possibly linker structures) between the single tubes^19^. N-Cadherin is a classical type I Cadherins that mediates Ca^2+^-dependent intercellular adhesion^20^. It engages in homophilic binding with N-Cadherin molecules on adjacent cells^21^ and interacts intracellularly with Cadherin-associated molecules including p120-Catenin, β-Catenin, and α-Catenin^22,23^. Among these, α-Catenin is crucial for anchoring the adhesion complex to the actin cytoskeleton, a requirement for its functional integrity^24,25^. α-Catenin is a versatile protein with multiple binding partners based on its conformational state^26,27^, therefore controlling actin dynamics. It can limit the formation of branched actin filaments by competing with Arp2/3 for actin binding^28^, induce the formation of filopodia by being recruited to phosphoinositide-activated membranes in the form of Cadherin-free homodimers^29^, or recruit formins involved in linear actin nucleation an elongation such as mDia1^30^. Based on previous data showing the presence of members of the Cadherin superfamily on TNTs^19,31–35^, we hypothesize that the Cadherin-Catenin complex plays a critical role in the structural regulation and functional dynamics of TNTs.

By combining quantitative assays in living cells with cryo-CLEM and tomography, here we demonstrate that N-Cadherin is a key organizer of both the structure and function of TNTs. In the absence of N-Cadherin the TNT bundle (formed by iTNTs) appears disorganized, unstable and nonfunctional. Conversely, N-Cadherin overexpression (OE) increases TNT stability and the transfer of vesicles within them. Our data further suggests that TNT-mediated transfer is facilitated by high cellular density and can occur even over short intercellular distances. However, the role of N-Cadherin in TNT regulation is mechanistically distinct from its effect on increasing cell density.

We also show that α-Catenin is required for TNT formation and acts downstream of N-Cadherin in this regulatory pathway. In addition, we identified roles for p120-Catenin (known to stabilize N-Cadherin at the membrane), as well as non-muscle myosin 2A (NMIIA) and its upstream regulator the Rho-associated kinase ROCK, as key contributors to TNT-mediated transfer, likely through modulation of cortical tension. We demonstrate the ability of NMIIA to both be processive inside TNTs and to transfer passively within using actin retrograde flow. Finally, we uncover a pathway involving the Rho GTPase Cdc42, its binding partner at the membrane IRSp53, and their downstream effector N-WASP to be essential in TNT-mediated transfer, possibly through the facilitation of membrane fusion.

## Results

### 1. N-Cadherin expression promotes contact-dependent transfer

N-Cadherin has previously been observed in TNTs in various cell types, including CAD, HeLa, urothelial, U2OS and HEK cells^19,31–35^, yet its functional role in TNT formation and TNT-mediated transfer remains unclear. To address this, we used SH-SY5Y human neuronal cells, a well-established model for various physiological and pathological neuronal contexts, previously characterized for both TNT formation and TNT-mediated cargo transport^7,11,14,35–38^.

Confocal immunofluorescence using an anti-N-Cadherin antibody revealed that, similar to CAD cells, N-Cadherin decorates TNT-like structures between SH-SY5Y cells (Fig. 1A). This localization was confirmed at the ultrastructural level using immunogold labeling and cryo-electron tomography (cryo-ET), which showed N-Cadherin on iTNT membranes and at sites of interconnections between iTNTs (Fig. 1B–D), suggesting a conserved ultrastructural organization of TNTs in neuronal cells of both human and mouse origin^19^.

**Fig. 1.**
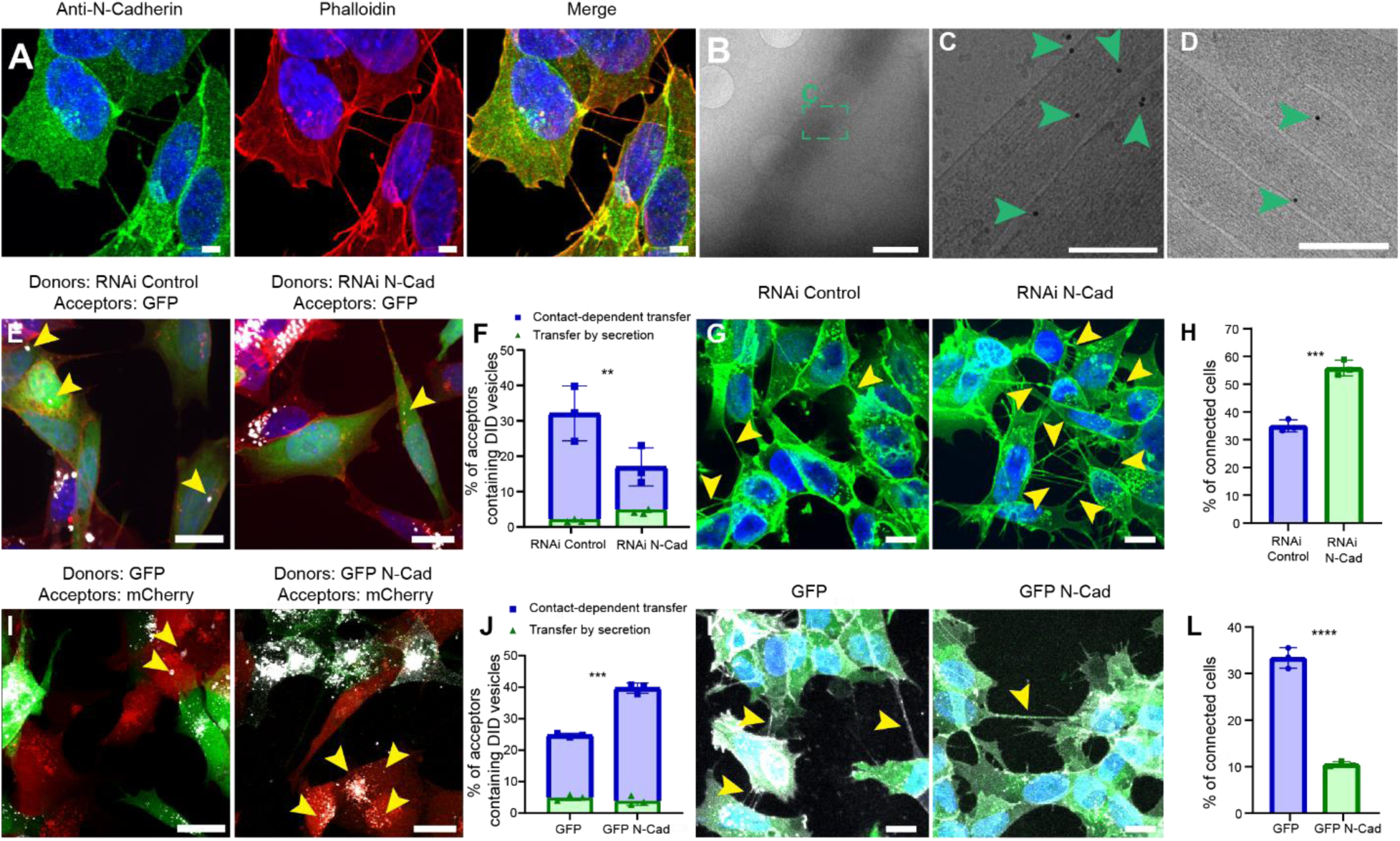
N-Cadherin localizes to TNT-like connections and modulates TNT functionality. (A) Immunofluorescence showing the localization of N-Cadherin along actin-based TNT-like connections. (B-C-D) Anti-N-cadherin immunogold labeling of TNT-like connections acquired by cryo-EM. (B) represents intermediate magnification. (C) High magnification cryo-tomography slices corresponding to the green square in (B). (D) High magnification cryo-tomography slices of iTNTs positive for N-Cadherin. Green arrows show N-cadherin targeted gold beads. (E). Representative confocal images showing 16h co-culture between (left) RNAi Control with DiD-labelled vesicles (donors) and GFP cells (acceptors), or (right) RNAi N-Cadherin (donors) challenged with DiD-labelled vesicles (white) and GFP cells (acceptors) (on the right). Cellular membranes were labelled with WGA-546 (red), nuclei were stained with DAPI (blue). The yellow arrowheads indicate DiD-labelled vesicles detected in the cytoplasm of acceptor cells. (F). Graph showing the percentage of acceptor cells containing DiD-labelled vesicles from the co-cultures in cells transfected with RNAi Control (32.12% ± 7.77) for contact-mediated transfer in blue, p; 2.31%± 1.31 for transfer by secretion in green) or RNAi N-Cadherin (17% ± 5.4 for contact-mediated transfer in blue; 5.03% ± 1.18 for transfer by secretion in green). (G) Confocal images of TNT-like connections between RNAi Control cells (on the left) RNAi N-Cadherin cells (on the right). Cells stained with WGA-488 (green) and DAPI (blue) for the nuclei. The yellow arrows indicate TNT-like connections. (H) Graph showing the percentage of connected cells transfected with RNAi Control non-targeting (35% ± 2.17) and RNAi N-Cadherin (55.8% ± 2.85). (I) Representative confocal images showing 16 h co-culture between DiD-labelled GFP cells (donor) and mCherry cells (acceptors) (on the left, GFP-N-Cadherin challenged with DiD-labelled vesicles (donor) and mCherry cells (acceptors) (on the right). The yellow arrowheads indicate DiD-labelled vesicles detected in the cytoplasm of acceptor cells. (J). Graph showing the percentage of acceptor cells containing DiD-labelled vesicles from the co-cultures in GFP control cells (24.78% ± 0.63 for contact-mediated transfer in blue; 4.97% ± 0.93 for transfer by secretion in green) against GFP N-Cadherin cells (39.71% ± 1.62 for contact-mediated transfer in blue; 3.95% ± 1.48 for transfer by secretion in green). (K) Confocal micrograph showing TNT-like connections between (left) GFP expressing cells or (right) between GFP N-Cadherin cells. Cells stained with WGA-647 (gray) and DAPI (blue) for the nuclei. The yellow arrows indicate the TNT-like connections. (L). Graph showing the percentage of connected cells in GFP expressing cells (33.4% ± 2.22) and GFP-N-Cadherin (10.5% ± 0.60). All experiments represent N=3. Data are presented as mean ± SD. Statistical significances were determined using an unpaired two-tailed Student’s t-test. *p < 0.05, **p < 0.01, ***p < 0.001. Scale bars in B = 2 μm, in C = 200 nm D=150 nm), and in E,G,I,K = 10 μm.

To assess N-Cadherin’s functional role, we measured both contact-mediated vesicle transfer and the proportion of TNT-connected cells following modulation of N-Cadherin expression. Acute knockdown (KD) of N-Cadherin via RNA interference (82% reduction compared to control; Fig. S1A) was followed by a vesicle transfer assay in co-culture^39^ (Fig. S1B, top). Donor cells containing DiD-labeled vesicles were co-cultured with acceptor cells expressing GFP (to distinguish them from donors) for 16 h. The proportion of acceptor cells positive for DiD-labeled vesicles was significantly lower in the N-Cadherin KD condition (17%) compared to the scramble siRNA control (32%) (Fig. 1E–F). To rule out secretion-mediated transfer, acceptor cells were incubated with conditioned media from control and KD donor cells (Fig. S1B, bottom), which resulted in minimal transfer (2.31% vs. 5.03%) and did not follow the same trend (Fig. 1F), supporting the conclusion that transfer occurs primarily through contact and is reduced by N-Cadherin knockdown.

Unexpectedly, quantification of TNT-like, non-adherent intercellular bridges showed an increased number of connected cells in the N-Cadherin KD condition (56%) compared to control (35%) (Fig. 1G–H).

To further probe the role of N-Cadherin, we generated a stable SH-SY5Y cell line overexpressing GFP-tagged N-Cadherin. The ectopically expressed protein exhibited a distribution similar to the endogenous one (Fig. S1C). When used as donors in co-culture, GFP-N-Cadherin cells transferred vesicles more efficiently (40%) than GFP-only controls (25%) (Fig. 1I–J). Again, secretion-based transfer remained minimal in both conditions (Fig. 1J), reinforcing a functional role of N-Cadherin in TNT-mediated transfer.

However, GFP N-Cadherin–expressing cells displayed a reduced percentage of TNT-like connections (10%) compared to GFP-only cells (33%) (Fig. 1K–L). Taken together, these findings reveal an inverse relationship between the number of TNT-like connections and the efficiency of contact mediated vesicle transfer. This is in contrast to earlier studies, which reported a direct correlation between the number of actin-based intercellular bridges and DiD-labeled vesicle transfer^39–41^.

A notable phenotype observed upon N-Cadherin overexpression (OE) was a significant increase in cell density (measured as the number of cells per µm^2^; see *Materials and Methods*), which could influence the morphological detection of TNT-like connections, as these are more easily visualized between well-separated cells^39–41^. Quantification confirmed a significant increase in cell density in N-Cadherin OE cells compared to controls (Fig. S1D), while RNAi-mediated knockdown significantly reduced density (Fig. S1E).

The area of isolated single cells remained unchanged in the N-Cadherin KD condition (Fig. S1G) but was significantly larger in the OE condition (Fig. S1F), possibly leading to an underestimation of local clustering in our density measurements. These findings suggest that the denser organization of N-Cadherin OE cells may reduce the detectability of individual TNT-like structures, which could partially explain the observed increase in contact-dependent DiD vesicle transfer.

### 2. TNT-mediated transfer is enhanced by N-Cadherin through both density-dependent and -independent mechanisms

Although TNTs are typically described as long structures given their capacity to exceed the length of canonical filopodia there is, to our knowledge, no data indicating that functional TNTs require distant cell positioning. On the contrary, we would expect shorter connections to form more readily between closely apposed cells, increasing the likelihood of material transfer. Therefore, the increased membrane apposition observed in N-Cadherin OE cells (Fig. S1F) may facilitate the formation of short, functional connections that evade detection using current morphological criteria^39^.

To test whether cell density affects TNT quantification, we plated SH-SY5Y cells at low (< 0.003 cell/μm^2^) and medium densities (< 0.0045 cell/μm^2^) (see *Material and Methods*) (Fig. 2A-C). Higher plating density significantly reduced the number of detectable TNT-like connections (Fig. 2D), supporting the idea that increased crowding impairs morphological identification.

**Fig. 2.**
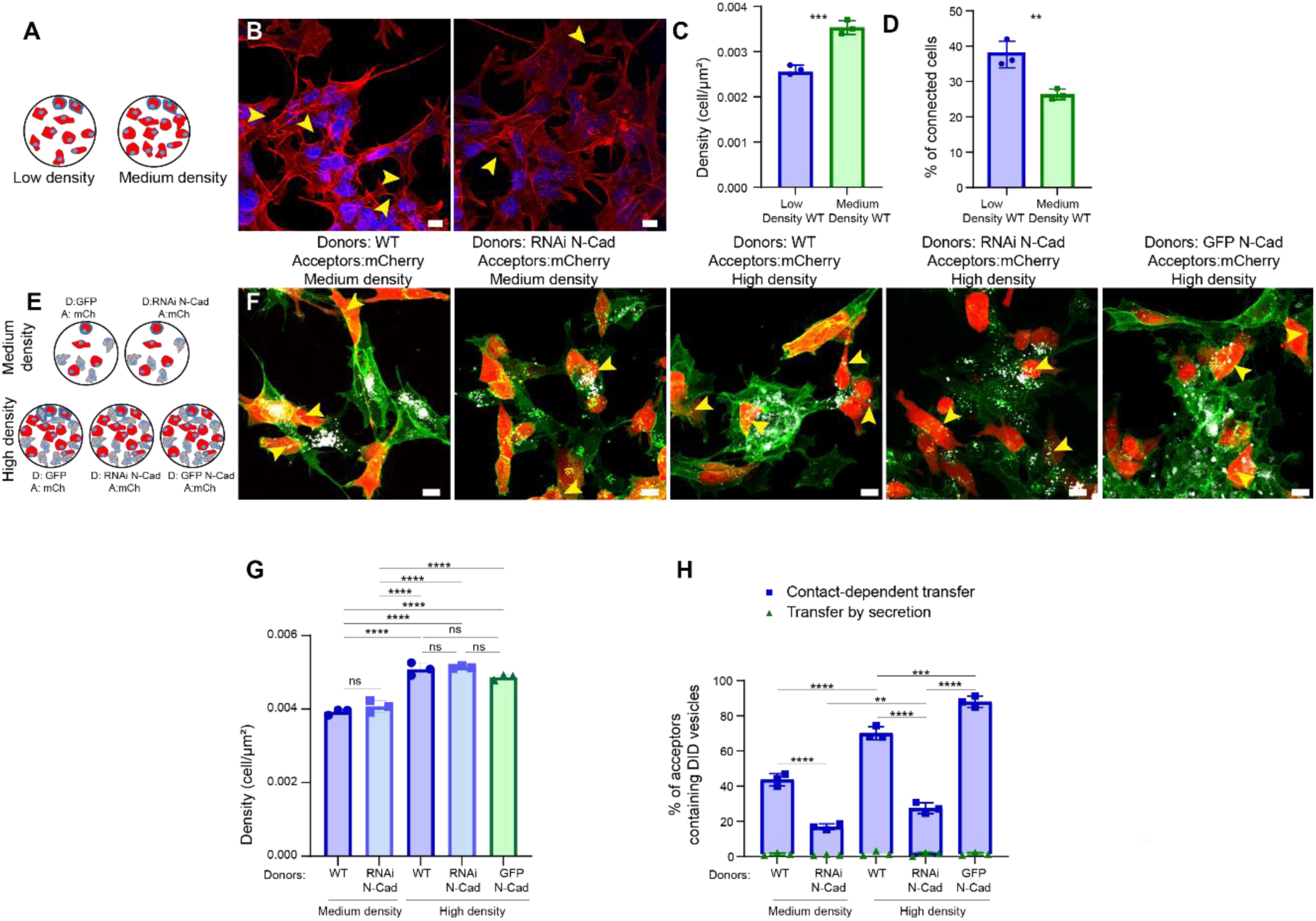
Higher cell density reduces observable TNT-like connections but enhances contact-dependent transfer, with N-cadherin further promoting transfer. (A) Schematic representing experimental design used for (B-D). (B) Graph showing the cell density of *Wild-Type* SH-SY5Y plated at low density (60 000 cells) (0.00260 ± 0.00010 cell/μm^2^) and medium density (100 000 cells, see *Material and Methods*) (0.00353 ± 0.00015 cell/μm^2^).Cell membranes were stained using WGA 546 (red) and DAPI (blue) for the nuclei. (C) Graph showing cell density of cells in (A-B) (from left to right, 0.00260 ± 0.00001, 0.00353 ± 0.00002 cell/μm^2^). (D) Graph showing the percentage of connected cells in low density *Wild-type* expressing cells from graph (B) (37.67% ± 3.79) and Medium density (26.33% ± 1.53). (E) Schematic representing experimental design used for (F-H). (F) Representative confocal images showing 16 h co-culture between various donor cells loaded with DiD-labelled vesicles (white) co-cultured at 1:1 ratio with mCherry cells (red) as acceptors plated at medium or high density. Cell membrane were stained using WGA 488 (green) for membrane. The yellow arrowheads indicate DiD-labelled vesicles detected in the cytoplasm of acceptor cells. (G) The graph shows the cell density measurement of cells represented in (D) (see *Material and Methods*). From left to right, in cell/μm^2^: 0.00393 ± 0.00008; 0.00407 ± 0.00016; 0.00508 ± 0.00017; 0.00513 ± 0.00004; 0.00487 ± 0.00007. (H) Graph showing the percentage of acceptor cells containing DiD-labelled vesicles from the co-cultures in GFP donor cells at medium density (45% ± 3, ∼1.2% via secretion), RNAi N-Cadherin GFP donor cells at medium density (17.67% ± 1.53, ∼0.5% via secretion), GFP donor cells at high density (71.67% ± 3.06, ∼0.6% via secretion), RNAi N-Cadherin GFP donor cells at high density (28.67% ± 3.79, ∼1.2% via secretion) or GFP-N-Cadherin donor cells at high density (89.33% ± 2.52, ∼0.8% via secretion). Green bars represent the percentages of DiD-positive acceptor cells in parallel secretion controls. All experiments represent N=3. Data are presented as mean ± SD. Statistical significances were determined using an unpaired two-tailed Student’s t-test for (C) and (D), and using a one-way ANOVA followed by Tukey’s post hoc test for (G) and (H). *p < 0.05, **p < 0.01, ***p < 0.001. Scale bars = 10 μm.

To explore whether close cell proximity might occult intercellular connections, we co-cultured cells overexpressing actin-Chromobody-RFP (aC-RFP) with cells overexpressing actin-Chromobody-GFP (aC-GFP). This allowed us to visualize non-substrate-adherent actin-rich protrusions that might otherwise be missed (Fig. S2A). As expected, short connections (< 5 μm) were substantially more abundant than long ones, suggesting that many TNTs may form between closely adjacent cells.

To unmask such connections, we exposed both control and GFP N-Cadherin OE cells to short (90-second) diluted trypsin pulses (see *Material and Methods*) to gently separate cell bodies while preserving intercellular structures^42^ (Fig. S2B). Under these conditions, the percentage of TNT-connected cells increased significantly in both control and OE cultures (Fig. S2C), with N-Cadherin OE cultures showing a significantly higher proportion of connected cells than control after trypsinization (78% vs. 67%).

These results indicate that by increasing cell density, N-Cadherin OE promotes the formation of short TNT-like structures that are less easily detected using standard morphological criteria^39^. As a result, the actual number of functional intercellular connections may be underestimated.

An important set of questions is whether the connections formed at higher cell densities are functional, that is, capable of supporting transfer, and whether N-Cadherin’s influence on TNTs is solely due to increased cell density or also involves additional mechanisms. To address this, we performed vesicle transfer assays under five conditions: (1) GFP donor cells plated at medium confluency, (2) RNAi N-Cadherin donor cells at medium confluency, (3) GFP-donor cells at high confluency, (4) RNAi N-Cadherin donor cells at high confluency, and (5) GFP N-Cadherin OE donor cells at high confluency. In each case, donor cells were co-cultured with mCherry OE acceptor cells at a 1:1 ratio (Fig. 2E-G). For each field of view, we measured cell density. Densities below 0.0045 cells/µm² were classified as “medium,” while values above this threshold were considered “high” for both mCherry acceptors and GFP N-Cadherin OE populations (Fig. 2G). Increasing cell density from medium to high in GFP donor cells led to a significant rise in contact-dependent transfer to mCherry acceptors (from 45% to 72%) (Fig. 2G–H). A similar trend was observed in RNAi N-Cadherin cells, with transfer increasing from 18% to 29% as density increased. These findings highlight the important, yet often overlooked, role of cell density in TNT-mediated transfer. However, even at comparable high density, N-Cadherin OE donor cells transferred significantly more vesicles than GFP controls (89% vs. 72%). Likewise, GFP donor cells at medium density transferred significantly more than RNAi N-Cadherin cells at high density (45% vs. 29%). These results suggest that N-Cadherin enhances TNT-mediated transfer through mechanisms at least partially independent of changes in cell density. As expected, secretion-mediated transfer remained negligible across all conditions. Based on these findings, we hypothesized that N-Cadherin might contribute to TNT stabilization and/or facilitate membrane fusion events required for the formation of functional, open-ended TNTs. To test TNT stability, we performed live-cell imaging to measure the lifetime of intercellular connections in N-Cadherin OE cells versus GFP controls (see *Material and Methods*) (Fig. S2D; Movies S1–S2). We found that TNTs in N-Cadherin OE cells were significantly more stable than those in controls (34,5 min vs 20 min). Together, these results indicate that N-Cadherin promotes efficient contact-dependent vesicle transfer not only by increasing cell clustering, which favors the formation of short, less detectable TNTs, but also by stabilizing these structures in accordance to what had been shown in HEK293 cells^34^, thereby enhancing their functionality.

### 3. N-Cadherin, α-Catenin, and p120-Catenin Cooperate in TNT-Mediated Transfer

To determine whether the effects on TNT formation and intercellular transfer are linked to the homotypic, transcellular binding function of N-Cadherin, we investigated the role of α-Catenin. This protein is essential for anchoring the Cadherin adhesion complex to the actin cytoskeleton, thereby reinforcing cell–cell adhesion through the Cadherin–actin “clutch” mechanism^43,44^.

We first analyzed α-Catenin expression and subcellular localization in our system (Fig. S3A). SH-SY5Y cells expressed endogenous α-Catenin localized at the plasma membrane, cytoplasm, and within TNT-like structures. Notably, α-Catenin co-localized extensively with N-Cadherin in these structures.

Next, we acutely knocked down α-Catenin in both donor and acceptor cells using RNAi, achieving an ∼85% reduction compared to RNAi controls (Fig. S3B). This led to a significant decrease in contact-mediated vesicle transfer, from 29% in controls to 18% in α-Catenin-depleted cells, while transfer via secretion remained minimal in both conditions (Fig. 3A-B). Interestingly, the percentage of TNT-connected cells increased upon α-Catenin knockdown (48% vs. 29% in controls; Fig. 3C-D), mirroring the phenotype observed with N-Cadherin interference.

**Fig. 3.**
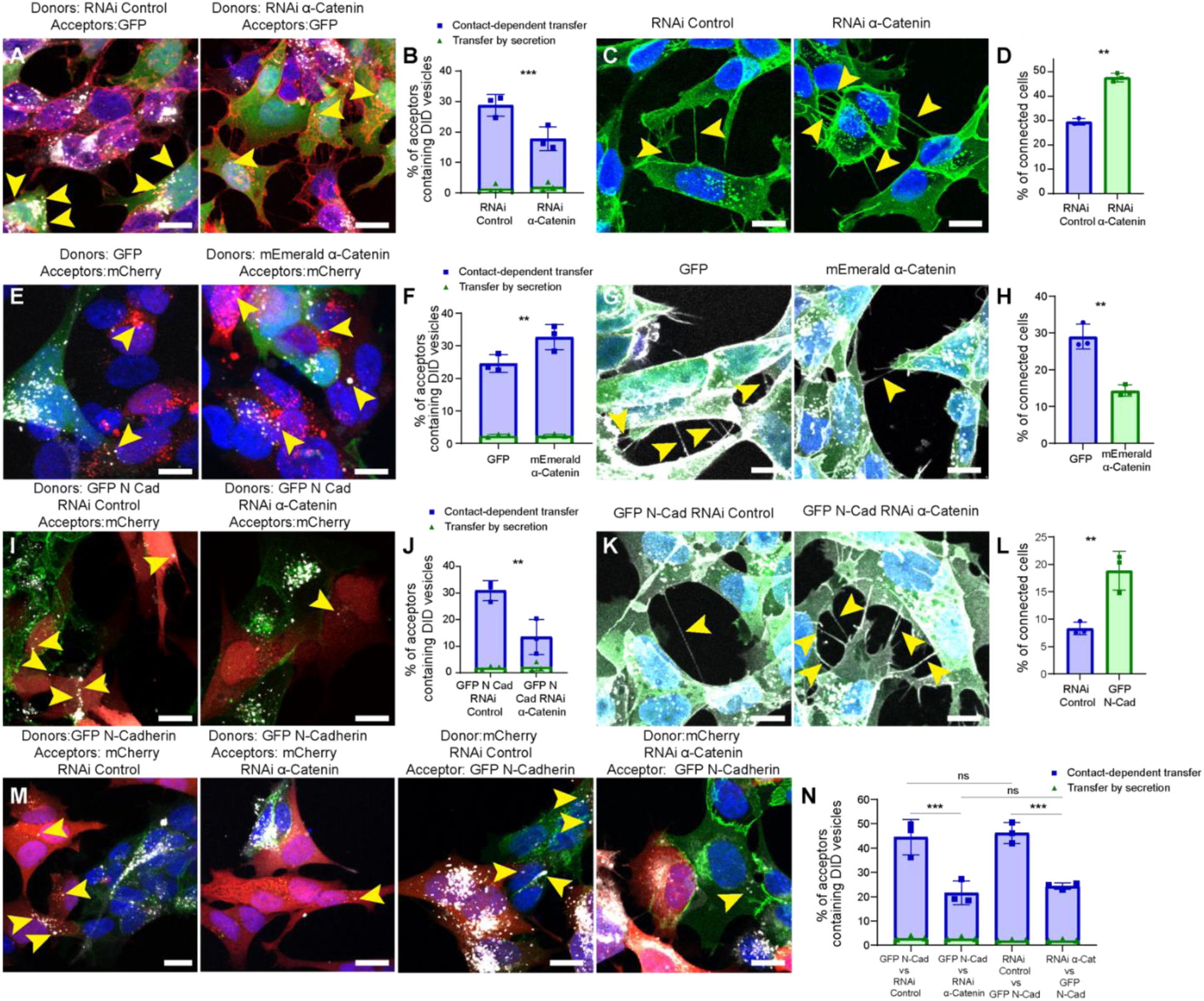
N-Cadherin-mediated promotion of TNT-functionality depends on the downstream interactor α-Catenin. (A) Representative confocal microscope images of an 16h co-culture of (left image) RNAi Control transfected donor cells challenged with DiD-labelled vesicles (white) and GFP acceptor cells (green) and (right image) RNAi α-Catenin transfected donor cells challenged with DiD-labelled vesicles (white) and GFP acceptor cells (green). Cellular membranes were labelled with WGA-546 (red) and nuclei were stained with DAPI (blue). The yellow arrowheads indicate DiD-labelled vesicles detected in the cytoplasm of acceptor cells. (B). Graph showing the percentage of acceptor cells containing DiD-labelled vesicles from the co-cultures in cells transfected with RNAi Control (28.74% ± 3.55 for contact-dependent transfer in blue; 1.5% ± 1.33 for transfer by secretion in green) or RNAi α-Catenin (17.75% ± 3.91 for contact-dependent transfer in blue; 2.08% ± 1.35 for transfer by secretion in green). (C). Representative confocal microscope images showing TNT-like structures between (left image) RNAi Control cells or (right image) RNAi α-Catenin cells. Cells stained with WGA-488 (green) and DAPI (blue) for the nuclei. The yellow arrowheads indicate the connections. (D). Graph showing the percentage of connected cells transfected with RNAi Control non-targeting (29.4% ± 1.31) and RNAi α-Catenin (47.6% ± 1.71). (E) Representative confocal microscope images of an 16h co-culture of (left image) GFP donors loaded with DiD-labelled vesicles (white) and mCherry cells acceptors (red) and (right image) mEmerald α-Catenin donors challenged with DiD-labelled vesicles (white) and mCherry acceptor cells (red). The yellow arrowheads indicate DiD-labelled vesicles detected in the cytoplasm of acceptor cells. (F). Graph showing the percentage of acceptor cells containing DiD-labelled vesicles from the co-cultures in GFP control cells (24.52% ± 2.70 for contact-mediated transfer in blue; 2.42% ± 0.42 for transfer by secretion in green) against mEmerald-α-Catenin cells (32.64% ± 3.90 for contact-mediated transfer in blue; 2.45% ± 0.39 for transfer by secretion in green). (G). Confocal micrograph showing TNT-like structures between (left) GFP cells or (right) mEmerald-α-Catenin cells. Cells stained with WGA-647 (grey) and DAPI (blue) for the nuclei. Yellow arrows indicate intercellular connections. (H) Graph showing the percentage of connected cells transfected with GFP (29% ± 3.38) and mEmerald-α-Catenin (14.3% ± 1.49). Cells were stained with WGA-647 (gray) and DAPI (blue) for the nuclei. (I) Representative confocal images of GFP N-Cadherin RNAi Control donors (green) with DiD-labelled vesicles in co-culture with mCherry acceptors (red) or (right) GFP N-Cadherin RNAi α-Catenin donors loaded with DiD-labelled vesicles (green) and mCherry acceptors (red). The yellow arrowheads indicate DiD-labelled vesicles detected in the cytoplasm of acceptor cells. (J) Graph showing the percentage of acceptor cells containing DiD-labelled vesicles from the co-cultures in GFP N-Cadherin cells transfected with RNAi Control (43.93% ± 2.59 for contact-mediated transfer in blue; 2.24%± 0.22 for transfer by secretion in green) or RNAi α-Catenin (23.53% ± 5.23 for contact-mediated transfer in blue; 2.36%± 1.78 for transfer by secretion in green). (K) Representative confocal images showing (left) TNT-like connections between GFP N-Cadherin cells with RNAi Control and (right) between GFP N-Cadherin cells with RNAi α-Catenin. Cells stained with WGA-647 (gray) and DAPI (blue) for the nuclei. The yellow arrows indicate the TNTs connected cells. (L). Graph showing the percentage of connected GFP N-Cadherin cells transfected with RNAi Control (8.32% ± 1.15) and RNAi α-Catenin (18.8% ± 3.54).(M) Representative confocal images of an 16h co-culture between (first image) GFP RNAi Control donors (green) with DiD-labelled vesicles (white) and mCherry cells acceptors (red) or (second image) GFP RNAi α-Catenin donors (green) challenged with DiD-labelled vesicles (white) and mCherry cells acceptors (red) or (third image) GFP N-Cadherin RNAi Control donors (green) with DiD-labelled vesicles (white) and mCherry cells acceptors (red) or (fourth image) GFP N-Cadherin RNAi α-Catenin donors (green) challenged with DiD-labelled vesicles (white) and mCherry cells acceptors (red). The yellow arrowheads indicate DiD-labelled vesicles detected in the cytoplasm of acceptor cells. (N) Graph showing the percentage of acceptor cells containing DiD-labelled vesicles from the co-cultures in GFP-N-Cadherin donor cells with RNAi Control acceptors (47.54%) or RNAi α-Catenin acceptors (24.49%) or GFP N-Cadherin acceptor cells with RNAi Control donors (48.34%) or RNAi α-Catenin donors (26.47%). Green bar represent parallel secretion controls. All experiments represent N=3. Data are presented as mean ± SD. Statistical significances were determined using an unpaired two-tailed Student’s t-test for (B, D, F, H, J, L), and using a one-way ANOVA followed by Tukey’s post hoc test for (N). Scale bars = 10 μm.

Conversely, overexpression of α-Catenin (mEmerald–α-Catenin) enhanced contact-mediated transfer, increasing DiD-positive acceptor cells from 25% to 33% (Fig. 4E-F). This increase in transfer, however, was accompanied by a notable reduction in TNT-connected cells (14% vs. 29% in controls; Fig. 4G-H). Thus, both knockdown and overexpression of α-Catenin recapitulated the effects of N-Cadherin perturbation (Fig. 1E, 1G), suggesting a cooperative role between the two in regulating TNT-mediated transfer.

**Fig. 4.**
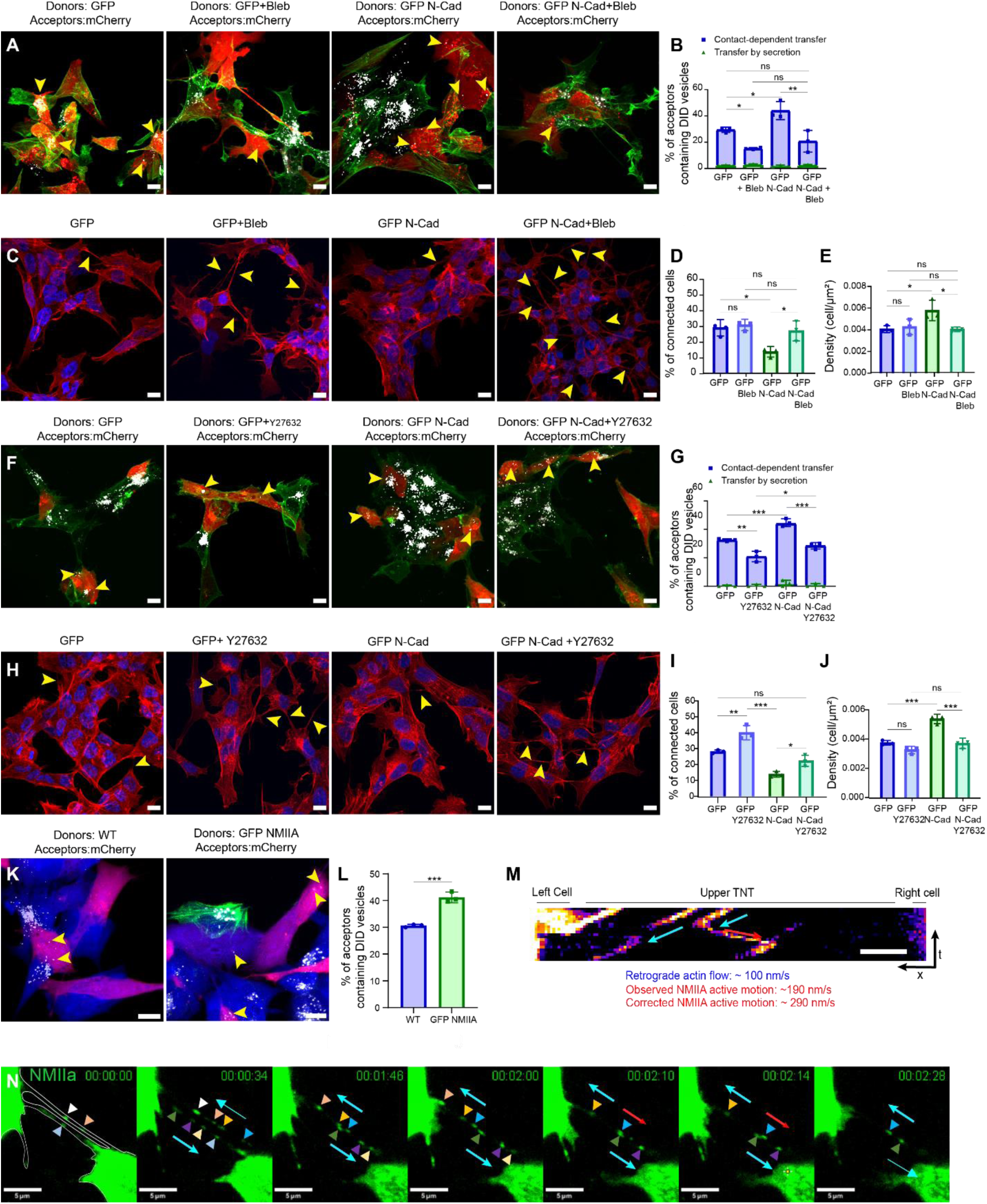
Effect of blebbistatin, Y-27632, and NMIIA expression on vesicle transfer and TNT connections in control and GFP N-Cadherin SH-SY5Y cells. (A) Confocal micrograph showing co-culture of donor SH-SY5Y cells (expressing GFP or GFP–N-Cad with DID 647) and acceptor SH-SY5Y cells (expressing mCherry), with or without blebbistatin treatment (10 μM). Yellow arrowheads indicate mCherry acceptors positive for DID vesicles. Cells stained with WGA-488 (green). (B) Graph showing the percentage of mCherry acceptor cells containing DiD-labelled vesicles from a co-culture with GFP or GFP-N-Cadherin donor cells with or without blebbistatin treatment (10 μM) (from left to right, 31.97% ± 1.94, 18.21% ± 1.08, 45.91% ± 6.38, 23.28% ± 7.95; secretion controls from left to right, 2.55% ± 0.17, 3.01% ± 0.49, 1.93% ± 0.63, 2.60% ± 0.39). (C) Confocal micrograph showing TNTs connected SH-SY5Y cells (expressing GFP or GFP–N-Cad), with or without blebbistatin treatment (10 μM). Cell membranes were stained using WGA 546 (red) and DAPI (blue) for the nuclei. The yellow arrows indicate the TNTs connected cells. (D) Graph showing the percentage of connected cells for GFP and GFP-N-Cadherin cells treated or not with blebbistatin (10 μM) (from left to right, 29.28% ± 5.21, 31% ± 3.61, 13.92% ± 3.42, 27.22% ± 6.37) (E) Graph showing cell density of cells in (C) (from left to right, 0.00406 ± 0.00032, 0.00428 ± 0.00071, 0.00579 ± 0.00094, 0.00401 ± 0.00021 cell/μm^2^). (F) Confocal micrograph showing co-culture of donor SH-SY5Y cells (expressing GFP or GFP–N-Cad upload with DID 647) and acceptor SH-SY5Y cells (expressing mCherry), with or without Y-27632 treatment (10 μM) Yellow arrowheads indicate mCherry acceptors positive for DID vesicles. Cells stained with WGA-488 (green). (G) Graph showing the percentage of mCherry acceptor cells containing DiD-labelled vesicles from a co-culture with GFP or GFP-N-Cadherin donor cells with or without Y-27632 treatment (10 μM) (from left to right, 32.54% ± 0.80, 21.17% ± 3.79, 46.84% ± 0.75, 29.16% ± 3.09; secretion from left to right, 0.3% ± 0.51, 0.5% ± 0.79, 2.1% ± 2.09, 0.7% ± 1.18). (H) Confocal micrograph showing TNTs connected SH-SY5Y cells (expressing GFP or GFP–N-Cad), with or without Y-27632 treatment (10 μM). Cell membranes were stained using WGA 546 (red) and DAPI (blue) for the nuclei. The yellow arrows indicate the TNTs connected cells. (I) Graph showing the percentage of connected cells for GFP and GFP-N-Cadherin cells treated or not with Y-27632 (10 μM) (from left to right, 28.08% ± 1.02, 39.99% ± 4.52, 13.84% ± 1.78, 22.33% ± 3.51). (J) Graph showing cell density of cells in (H) (from left to right, 0.00374 ± 0.00016, 0.00325 ± 0.00026, 0.00537 ± 0.00032, 0.00370 ± 0.00036 cell/μm^2^). (K) Confocal micrograph showing co-culture of donor SH-SY5Y cells control and expressing GFP NMIIA expressing. Cell cytoplasm was stained using Cell Mask blue. Yellow arrowheads indicate mCherry acceptors positive for DID vesicles (L) Graph showing the percentage of mCherry acceptor cells containing DiD-labelled vesicles from a co-culture with WT or GFP NMIIA donor cells (10,60 ± 1,171). (M) Kymograph of GFP-NMIIA filaments motion inside the upper TNT of the movie in N, from 1:48 min to 2:26 min. (N) Movie panel of GFP-NMIIA transfer within TNTs of opposite orientations connecting two SH-SY5Y cells. Movie Scale bars, 10 μm. All experiments represent N=3. Data are presented as mean ± SD. Statistical significances were determined using an unpaired two-tailed Student’s t-test for (L), and using a one-way ANOVA followed by Tukey’s post hoc test for (B, D, E, G, I, J). *p < 0.05, **p < 0.01, ***p < 0.001.

To assess whether α-Catenin acts downstream of N-Cadherin, we performed KD and OE experiments in N-Cadherin-overexpressing cells. In GFP N-Cadherin and mCherry-expressing cells, α-Catenin knockdown reduced protein levels by 74% and 80%, respectively, compared to RNAi controls (Fig. S3C). Importantly, N-Cadherin levels and localization remained unaffected (Fig. S3D). Functionally, α-Catenin depletion in GFP N-Cadherin OE donor cells led to a significant decrease in contact-mediated transfer (from ∼44% to 24%), while secretion-mediated transfer remaining minimal (Fig. 3I-J). Additionally, TNT-like connections were more frequent in GFP N-Cadherin/α-Catenin KD cells compared to controls (19% vs. 8%; Fig. 3K-L). These findings indicate that α-Catenin depletion overrides the effects of N-Cadherin OE, supporting its role as a key downstream effector in contact-dependent transfer.

Because cadherins-based adhesion depends on mechanical coupling between opposing cortices, we hypothesized that N-Cadherin–dependent transfer would be impaired if α-Catenin was depleted in either the donor or acceptor cells. To test this, we performed DiD-vesicle transfer assays under two conditions: (i) GFP N-Cadherin donor cells co-cultured with either RNAi control or α-Catenin KD acceptors; and (ii) RNAi control or α-Catenin KD donor cells co-cultured with GFP N-Cadherin acceptors (Fig. 3M-N). In control conditions, contact-mediated transfer occurred at ∼44% (donor GFP N-Cadherin) and 46% (acceptor GFP N-Cadherin). However, α-Catenin knockdown in either population halved the transfer rate to ∼22–24% (Fig. 3N). Secretion-based transfer remained minimal in all conditions (Fig. 3N). These results indicate that α-Catenin is required in both donor and acceptor cells for efficient N-Cadherin– dependent transfer.

We next investigated the role of p120-Catenin, another member of the N-Cadherin-α-Catenin complex known to regulate cortical tension and stabilize N-Cadherin at the membrane ^45^. RNAi-mediated knockdown of p120-Catenin (average reduction of ∼83%) (Fig. S3F) significantly reduced contact-mediated DiD transfer (from 31.1% to 21.6%; Fig. S3G) and increased the percentage of TNT-connected cells (from 30.7% to 48.7%; Fig. S3H). This phenotype closely resembles that of N-Cadherin and α-Catenin knockdown, suggesting that p120-Catenin promotes TNT-mediated transfer by stabilizing N-Cadherin at cell-cell contacts.

Together, these findings strongly suggest that N-Cadherin, α-Catenin, and p120-Catenin function together to promote TNT-mediated intercellular transfer by positively regulating N-Cadherin homotypic binding.

### 4. NMII and ROCK regulate N-cadherin binding and TNT-Mediated Transfer, with NMIIA actively walking inside TNTs and transferring through them via the actin retrograde flow

Beacuse both α-Catenin and p120-Catenin regulate cortical tension, a key modulator of Cadherin-mediated adhesion^43–46^, we tested whether disrupting acto-myosin contractility would affect transfer efficiency. In myoblasts, inhibition of non-muscle myosin II (NMII) with blebbistatin reduces N-Cadherin’s accumulation at cell-cell junctions^47^. Similarly, Y-27632-mediated inhibition of ROCK, an upstream regulator of NMII’s contractility^48^, produces comparable effects^47^.

We treated control and N-Cadherin OE cells with blebbistatin (10 μM) and measured both transfer of DiD-labelled vesicles and intercellular connections. Blebbistatin significantly reduced the proportion of DiD-positive acceptor cells (from 32% to 18% in controls; from 46% to 23% in N-Cadherin OE cells; Fig. 4A-B). Interestingly, the percentage of TNT-connected cells was unchanged in control cells (29% vs. 28%) but increased upon blebbistatin addition in N-Cadherin OE cells following treatment (14% to 27%; Fig. 4C-D). While blebbistatin did not alter cell density in control cells, it significantly reduced density in N-Cadherin OE cultures (Fig. 4F), indicating that NMII activity contributes to the N-Cadherin–induced cell aggregation and impacts TNT quantification.

To further validate these findings, we inhibited ROCK with Y-27632 (10 μM, 18 hours) before fixing the cells. ROCK inhibition mirrored the effects of blebbistatin, significantly reducing transfer in both Control (from 36% to 21%) and N-Cadherin OE cells (from 47% to 29%; Fig. 4F-G). In addition, it increased the percentage of connected cells in control cells (from 28% to 40%) and in N-Cadherin OE cells (from 14% to 23%; Fig. 4H-I). Yet we only observed a significant reduction in cell density in N-Cadherin OE cells (Fig. 4J), highlighting both the importance of ROCK for stable N-Cadherin engagement and a possible density-independent role for ROCK in protrusion formation.

This supports a model in which N-Cadherin-mediated adhesion, modulated by cortical tension, enhances intercellular transfer through mechanisms that are at least partly independent of cell density.

NMIIA, but not NMIIB, has been shown to be present at the base of Myosin X-induced filopodia and promote their stabilization and elongation^49^, Myosin X having also been shown to promote the formation of TNTs in mouse neurons^40^. In order to better understand the role of NMIIA in TNT-mediated transfer, we cocultured mCherry acceptors with DiD-challenged donor cells transfected or not with GFP-NMIIA (Fig. 4K). We observed a significant increase in DiD-positive acceptors upon NMIIA OE (from 30.6% to 41.2%) (Fig. 4L).

NMIIA are non-processive as monomers, but can bundle into NMIIA filaments which have recently been shown to be capable of processive motility^50^. To assess NMIIA localization and motility inside TNTs, we performed fast live imaging of GFP-NMIIA upon OE in SH-SY5Y cells. We observed some processive NMIIA filaments motion inside TNTs. Addition, NMIIA filaments transfer between cells through TNTs passively, using the retrograde flow of actin (Fig. 4M-N, Movie S3).

This data demonstrates the importance of NMII and ROCK activity for TNT-mediated transfer in our cells, especially upon N-Cadherin overexpression. Furthermore, it hints at a transfer mechanism inside iTNTs dependent on F-actin retrograde flow rather than active motor proteins. Together, these findings demonstrate that N-Cadherin, α-Catenin, and p120-Catenin function together to promote TNT-mediated intercellular transfer. This process is finely regulated by cortical tension via NMII and ROCK activity and requires coordinated cadherin– actin coupling in both donor and acceptor cells.

### 5. N-Cadherin/α-Catenin Complex Regulates TNT Ultrastructure

N-Cadherin may stabilize tunneling nanotubes (TNTs) through two complementary mechanisms: (i) cross-linking individual TNTs (iTNTs) into parallel bundles that provide structural support and (ii) reinforcing TNT tip adhesion to recipient cells to facilitate docking and fusion. To investigate how N-Cadherin influences TNT architecture, we used cryo-electron tomography (cryo-ET) to analyze TNT ultrastructure in N-Cadherin knockdown (KD) cells.

Due to cryo-EM limitations, such as poor visibility in dense cellular areas and difficulty resolving short intercellular projections, we focused on long, actin-positive TNTs. In control SH-SY5Y cells, TNTs typically appeared as bundles of 2–6 parallel iTNTs (Fig. 5A–B), consistent with previous reports in mouse CAD neuronal cells^19^.

**Fig. 5.**
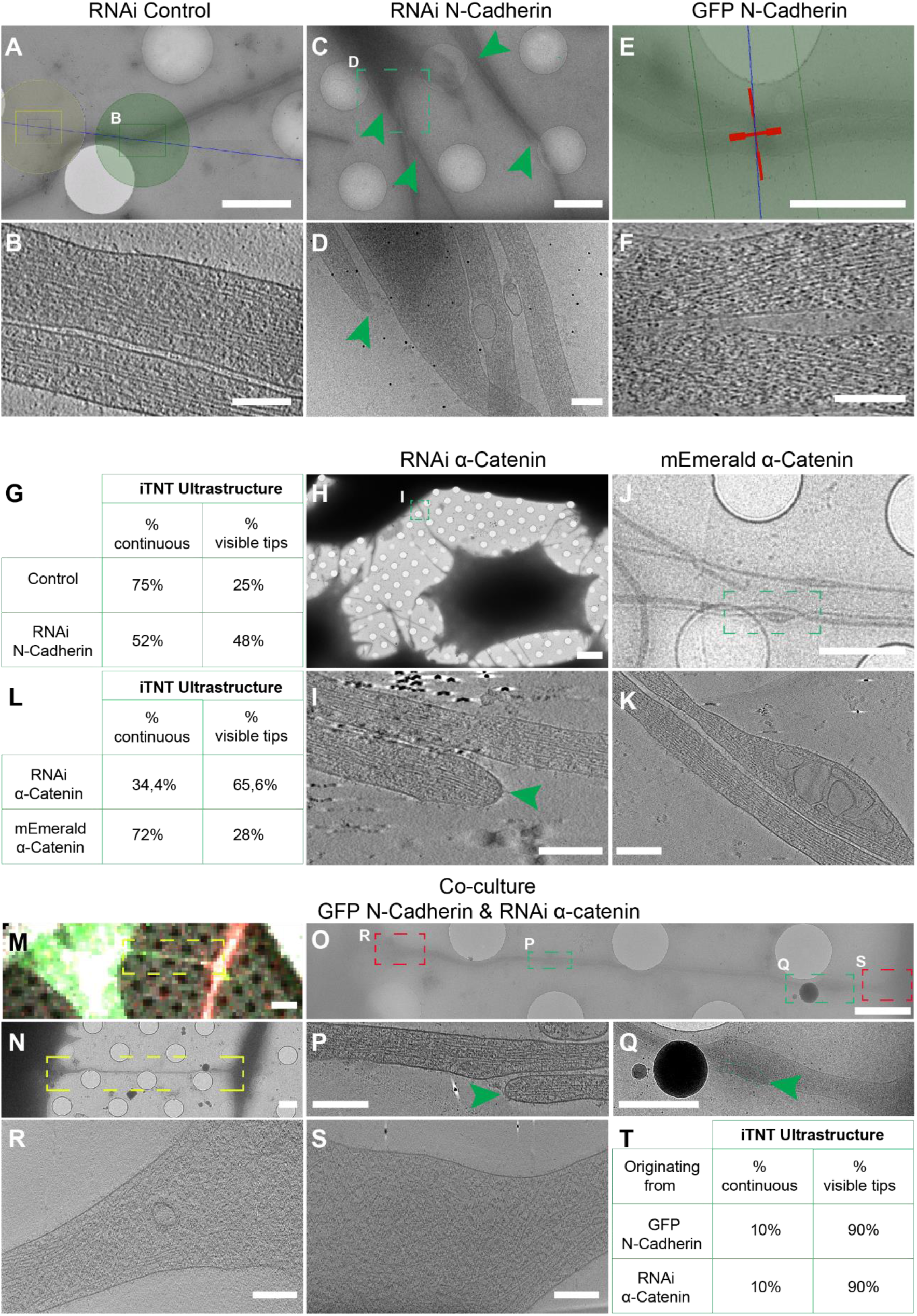
Cryo-EM of TNTs Formed Between SH-SY5Y Cells upon Up- or Downregulation of N-Cadherin and α-Catenin. (A). Cryo-EM intermedia micrograph showing TNT-connected RNAi Control cells. (B). High-magnification cryo-tomography slices corresponding to the green dashed squares in (A) showing continuous iTNTs. (C-D). TNT-connected RNAi N-Cadherin cells acquired by cryo-EM (C) low magnification. (D). High-magnification cryo-EM slices showing the iTNT in the green dashed square in (C). (E-F). TNT-connected GFP N-Cadherin cells acquired by cryo-EM (E) low magnification. (F). High-magnification cryo-tomography slices corresponding to the red cross in (E). (G). Table showing the percentage of continuous iTNTs and visible tips iTNTs. (H-I). Cryo-EM grids were prepared using RNAi α-Catenin cells (H) Low magnification of cryo-EM micrograph showing TNTs connecting RNAi α-Catenin cells. (I) High-magnification cryo-tomography slices of the dashed square in (H) showing iTNTs with visible tips (J-K). Cryo-EM grids were prepared using mEmerald α-Catenin cells. Low (J) magnification of cryo-EM micrograph showing TNT connecting mEmerald α-Catenin cells. (K) High-magnification cryo-tomography slices of the yellow dashed square in (J) showing parallel and continuous iTNTs. (L) Table showing the percentage of continuous iTNTs and visible tips iTNTs in cells downregulated for or overexpressing α-Catenin. (M). Confocal micrograph showing TNTs between RNAi α-Catenin (mCherry) and N-Cadherin (GFP) cells plated on EM grids. (N). Low cryo-EM micrograph showing TNT-connected cells in the dashed yellow square in (M). (O). Intermedia cryo-EM micrograph of (M). (P). High-magnification cryo-tomography slices corresponding to the green dashed square in (N). (P) Intermedia cryo-EM micrograph corresponding to the green dashed square in (O). (R). High-magnification cryo-tomography slices corresponding to the red dashed square in (O). (S). High-magnification cryo-tomography slices corresponding to the red dashed square in (O). (T). Table showing the percentage of TNTs with continuous and visible tips in between RNAi α-Catenin (mCherry) and GFP N-Cadherin, depending on the cell they originate from. Green arrowheads indicate visible tips iTNTs. Scale bars (A, C, E, M, N, J) 2 μm; (B, D, F, P, R, S,) 100 nm; (H) 10 μm; (K) 50 nm; (I) 100 nm.

In contrast, N-Cadherin KD cells exhibited disorganized, braided iTNTs with a marked increase in closed-ended protrusions (Fig. 5C–D, G). In N-Cadherin OE cells, as expected from increased cell clustering, long TNTs were more difficult to isolate, but where visible, iTNTs retained a parallel organization (Fig. 5E–F).

Quantitative analysis of TNT morphology revealed that in control cells, 75% of TNTs were fully extended between cells, while 25% had closed tips. In N-Cadherin KD cells, only 52% of TNTs were continuous, with 48% displaying closed ends (Fig. 5G). This increase in closed-ended structures suggests a failure of TNTs to dock or fuse with target cells in the absence of N-Cadherin. However, we note that cryo-EM resolution may not allow reliable discrimination of attachment points due to cell body thickness (>500 nm), limiting precise classification of open versus closed TNTs. Nonetheless, these structural changes align with the reduced intercellular transfer observed in N-Cadherin KD cells (Fig. 1E).

Given the physical association between N-Cadherin and α-Catenin, and to complement our confocal experiments (see above), we next assessed whether perturbing α-Catenin would induce similar ultrastructural changes. In α-Catenin KD cells, TNTs lost their parallel organization and instead displayed braided morphologies with a higher incidence of closed tips (Fig. 5H–I). In contrast, TNTs in mEmerald–α-Catenin OE cells remained predominantly parallel, resembling those in wild-type cells (Fig. 5J–K). Quantification confirmed these observations: in α-Catenin KD cells, only 35% of iTNTs were fully extended, while 65% had closed tips. In α-Catenin OE cells, 72% were extended and 28% were closed (Fig. 5L).

To determine whether α-Catenin is required in both donor and recipient cells, we performed correlative cryo-TEM on co-cultures of N-Cadherin OE cells with α-Catenin KD cells. We specifically selected TNTs for which we could unambiguously identify the donor and acceptor origin and assess tip morphology. Strikingly, in these mixed cultures, TNTs contained iTNTs from both cell types but nearly all showed closed ends, regardless of origin (Fig. 5M–S). Quantification showed that 90% of iTNTs in these heterotypic interactions were closed-ended (Fig. 5T), indicating that α-Catenin is essential in both donor and recipient cells for the formation of functionally open TNTs. This finding is consistent with our transfer assays, in which α-Catenin KD in either cell population reduced transfer efficiency even in the presence of N-Cadherin overexpression (Fig. 3N).

Together, these data indicate that the N-Cadherin/α-Catenin complex is critical for maintaining TNT ultrastructure and ensuring successful tip fusion, thereby supporting both the architectural stability and functional connectivity of TNTs. Furthermore, the requirement for bidirectional expression underscores the role of homotypic interactions in mediating TNT formation and intercellular communication.

### 6. N-Cadherin Overexpression Upregulates Small Rho GTPases, that Promote TNT-Mediated Transfer

To further elucidate the mechanism by which N-Cadherin promotes TNT-mediated transfer, we investigated the role of the small Rho GTPase Cdc42, a key regulator of actin remodeling and Cadherin-mediated adhesion. In macrophages, Cdc42 inhibition, as well as mutations in its effectors WASP and WAVE2, impairs the formation of TNT-like structures, with WASP knockdown also affecting TNT functionality^51^.

We first examined how N-Cadherin levels influence Cdc42 expression. Western blot analysis revealed that Cdc42 protein levels were significantly elevated in GFP N-Cadherin overexpressing cells, and conversely reduced two-fold in N-Cadherin knockdown (KD) cells compared to their respective controls (Fig. S4A–B). While these results do not directly address membrane-localized activity, they suggest that N-Cadherin modulates Cdc42 expression.

To assess the functional relevance of Cdc42, we performed DiD-based vesicle transfer assays where GFP or GFP N-Cadherin OE donor cells were treated or not with the Cdc42 inhibitor ML141 (5 μM). In GFP control cells, Cdc42 inhibition significantly reduced DiD-positive acceptor cells from 36% to 16%. The effect was even more pronounced in N-Cadherin OE cells, where transfer dropped from 53% to 19% (Fig. 6A-B). To demonstrate the robustness of our secretion control, we controlled for secretion using ThinCerts, where donors were cultured on top of acceptors cells, separated by a membrane of 1μm pore size (Fig. S1B, right, see *Materials and Methods*). Secretion-mediated transfer remained minimal across conditions, reflecting impaired contact-dependent transfer.

**Fig. 6.**
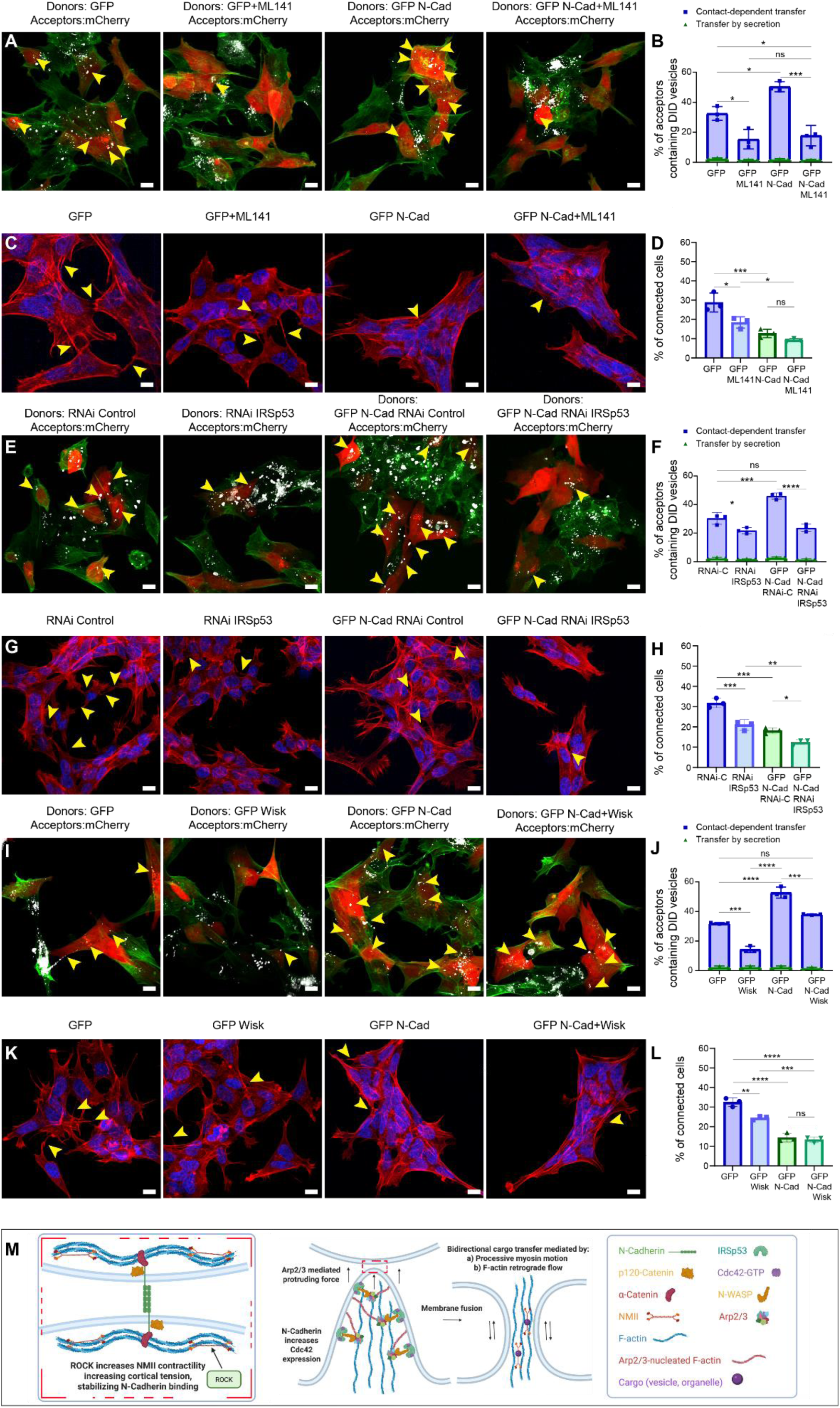
N-Cadherin expression positively correlates with Cdc42 expression, which promotes contact-dependent transfer along with downstream effectors IRSp53 and N-WASP. (A) Confocal micrograph showing co-culture of donor SH-SY5Y cells (expressing GFP or GFP–N-Cad with DID 647) and acceptor SH-SY5Y cells (expressing mCherry), with or without ML141 treatment (5 μM). Cells stained with WGA-488 (green). Yellow arrowheads indicate mCherry acceptors positive for DID vesicles. (B) Graph showing the percentage of mCherry acceptor cells containing DiD-labelled vesicles from a co-culture with GFP or GFP-N-Cadherin donor cells with or without ML141 treatment (5 μM) (from left to right, 35.02% ± 4.63, 16.97% ± 6.42, 52.25% ± 3.92, 19.26% ± 6.56; secretion controls from left to right, 2.4% ± 0.58, 1.5% ± 0.35, 1.8% ± 0.67, 1.4% ± 0.46). (C) Confocal micrograph showing TNTs connected SH-SY5Y cells (expressing GFP or GFP–N-Cad), with or without ML141 treatment (5 μM) Cell membranes were stained using WGA 546 (red) and DAPI (blue) for the nuclei. The yellow arrows indicate the TNTs connected cells. (D) Graph showing the percentage of connected cells for GFP and GFP-N-Cadherin cells treated or not with ML141 (5 μM) (from left to right, 28.79% ± 4.94, 18.38% ± 2.94, 12.73% ± 2.17, 9.30% ± 0.75). (E) Confocal micrograph showing co-culture of donor SH-SY5Y cells upon IRSp53 KD in GFP and GFP-N-Cadherin cells and acceptor SH-SY5Y cells (expressing mCherry). Cells stained with WGA-488 (green). Yellow arrowheads indicate mCherry acceptors positive for DID vesicles. (F) Graph showing the percentage of mCherry acceptor cells containing DiD-labelled vesicles from a co-culture with GFP or GFP-N-Cadherin donor cells transfected with either RNAi Control or RNAi IRSp53 (from left to right, 32.87% ± 3.55, 23.70% ± 2.13, 48.85% ± 2.64, 25.23% ± 2.09; secretion controls from left to right, 2.47% ± 0.70, 1.90% ± 0.35, 2.9% ± 0.79, 1.6% ± 0.57). (G) Confocal micrograph showing TNTs connecting cells upon IRSp53 KD in GFP and GFP-N-Cadherin cells. Cell membranes were stained using WGA 546 (red) and DAPI (blue) for the nuclei. The yellow arrows indicate the TNTs connected cells. (H) Graph showing the percentage of connected cells for GFP and GFP-N-Cadherin cells transfected with either RNAi Control or RNAi IRSp53 (from left to right, 31.26% ± 2.42, 22.83% ± 2.61, 15.60% ± 3.1, 12.53% ± 2.34). (I) Confocal micrograph showing co-culture of donor SH-SY5Y cells (expressing GFP or GFP–N-Cad with DID 647) and acceptor SH-SY5Y cells (expressing mCherry), with or without wiskostatin treatment (10 μM). Cells stained with WGA-488 (green). Yellow arrowheads indicate mCherry acceptors positive for DID vesicles. (J) Graph showing the percentage of connected cells for GFP and GFP-N-Cadherin cells treated or not with wiskostatin (10 μM) (from left to right, 32.49% ± 2.17, 24.50% ± 0.99, 14.47% ± 2.20, 13.40% ± 1.45). (K) Confocal micrograph showing TNTs connected SH-SY5Y cells (expressing GFP or GFP–N-Cad), with or without ML141 treatment (5 μM). Cell membranes were stained using WGA 546 (red) and DAPI (blue) for the nuclei. The yellow arrows indicate the TNTs connected cells. (L) Graph showing the percentage of connected cells for GFP and GFP-N-Cadherin cells treated or not with wiskostatin (10 μM) (from left to right, 32.49% ± 2.17, 24.50% ± 0.99, 14.47% ± 2.20, 13.40% ± 1.45). (M) Schematic of the membrane fusion model during TNT formation. Cdc42, upregulated by N-Cadherin, binds IRSp53 in highly curved protruding membranes, recruiting N-WASP itself recruiting Arp2/3. Local Arp2/3-mediated actin polymerization then promotes membrane fusion by exerting a critical force. All experiments represent N=3. Data are presented as mean ± SD. Statistical significances were determined using a one-way ANOVA followed by Tukey’s post hoc test for (B, D, F, H, J, L). *p < 0.05, **p < 0.01, ***p < 0.001.

To directly link these effects to TNT formation, we quantified TNT-connected cells. Cdc42 inhibition significantly reduced the proportion of connected cells in GFP controls (29% to 19%), but not in N-Cadherin OE cells (Fig. 6C-D). Interestingly, cell density increased modestly in both groups (significantly in N-Cadherin OE; Fig. S4C), suggesting that the observed drop in transfer was not due to reduced cell contact but rather to impaired TNT formation or stabilization.

Since Cdc42 cooperates with IRSp53 to recruit downstream actin modulators like N-WASP and Arp2/3, we next examined whether IRSp53 mediates Cadherin-dependent TNT transfer. RNAi knockdown of IRSp53 (56% and 76% reduction in control and N-Cadherin OE cells, respectively; Fig. S4D-E) significantly decreased transfer from 32% to 23% in control cells and from 49% to 23% in N-Cadherin OE cells (Fig. 6E-F). This abolishment of the N-Cadherin transfer gain suggests IRSp53 is required for Cadherin-enhanced transfer.

Consistent with this, IRSp53 knockdown reduced TNT connectivity (from 33% to 22% in controls, and 22% to 16% in N-Cadherin OE; Fig. 6G-H), with no significant changes in cell density (Fig. S4F), indicating that IRSp53’s role is independent of adhesion or cell clustering.

Finally, we inhibited N-WASP with Wiskostatin, observing reduced transfer in both control (33% to 16%) and N-Cadherin OE cells (54% to 40%) (Fig. 6I-J). TNT connectivity decreased only in control cells (from 31% to 24%), while remaining unchanged in N-Cadherin OE cells (13% vs. 12%) (Fig. 6K-L). Again, cell density was unaffected (Fig S4G), reinforcing the conclusion that this pathway acts independently of N-Cadherin-induced aggregation.

Together, these findings support the involvement of a Cdc42–IRSp53–N-WASP axis downstream of N-Cadherin in promoting TNT-mediated intercellular transfer. This mechanism is distinct from the increased transfer observed under high-density conditions, highlighting that N-Cadherin regulates both mechanical adhesion and actin-driven protrusive signaling to facilitate TNT function.

## Discussion

The findings of this study provide compelling evidence that N-Cadherin promotes contact-dependent intercellular transfer via TNTs. We demonstrate that increased cell density, regulated in part by N-Cadherin, enhances transfer, supporting previous observations that Cadherin-mediated adhesion promotes membrane apposition, a prerequisite for membrane fusion. This is consistent with prior work in epithelial cells, where N-Cadherin downregulation disrupted syncytium formation in the context of viral fusogen expression⁵⁰.

Interestingly, we also observed that higher cell density correlates with a reduced percentage of detectable TNT-like connections. However, this likely reflects an underestimation, due to technical limitations in detecting short connections between tightly packed cells, particularly under fixed or low-resolution imaging conditions.

Our data indicate that short connections form abundantly between cells in close contact. This underscores a broader challenge in TNT research: the lack of definitive markers and the reliance on structural features, such as length, to distinguish TNTs from other actin-based protrusions. While long TNTs are traditionally linked to material transfer due to their visibility, the high prevalence of short intercellular connections suggests that much of the transfer may occur through these less detectable pathways. However, capturing DiD-labelled vesicle transfer through short connections using live imaging proved technically challenging, primarily due to the high temporal resolution needed to track fast-moving particles within the cell volume. This limitation likely contributes to the frequent observation of TNT-mediated transfer in long connections rather than short ones. Nevertheless, given the numerical dominance of short intercellular structures, it is likely that TNT-mediated transfer occurs more frequently through these connections, both in our model and in other systems ^38,52,53^.

This insight calls for a reevaluation of our understanding of TNT-mediated transfer, especially in the context of dense tissues, where such short-range communication may be more relevant. Notably, a recent study using rapid live imaging identified fast, repeated vesicle exchanges between adjacent cells at specific membrane sites^54^. While the mechanism was not fully characterized, it is conceivable that short TNTs play a role in such exchanges.

Further supporting a role for N-Cadherin in membrane fusion, we observed an accumulation of closed TNT tips following N-Cadherin or α-Catenin knockdown, suggestive of incomplete fusion events. Cryo-ET further revealed that N-Cadherin is essential for maintaining the structural and functional integrity of TNT-like connections; N-Cadherin RNAi disrupted their canonical architecture, typically composed of parallel bundles of iTNTs. In contrast, N-Cadherin overexpression resulted in highly organized, tightly bundled iTNTs extending toward neighboring cells. The observed braiding of N-Cadherin-cross-linked iTNTs may result from NMII activity, as previously suggested in closed-ended TNTs in HeLa cells^33^. Reduced Cadherin-mediated adhesion could enhance braiding, whereas N-Cadherin overexpression may stabilize these structures, mechanically opposing myosin-driven contractility. This provides a mechanistic explanation for how adhesion molecules can influence the geometry and function of long-range cellular protrusions.

Importantly, our results show that the effect of N-Cadherin on TNTs cannot be attributed solely to increased cell density. Using a combination of KD, OE, and pharmacological inhibition in transfer assays alongside correlative cryo-EM imaging, we demonstrate that N-Cadherin’s role in TNT functionality is regulated by a network of signaling and structural proteins. These include p120-Catenin and α-Catenin, core components of the Cadherin adhesion complex well as regulators of cortical tension such as ROCK and NMII. The importance of cortical tension in Cadherin-mediated intercellular adhesion has been well documented^43,46^. We found that inhibition of ROCK or NMII led to a more pronounced reduction in transfer in N-Cadherin OE cells compared to controls, suggesting that these factors modulate contact-dependent transfer via N-Cadherin. Furthermore, inhibition of ROCK and NMII significantly reduced cell density in N-Cadherin OE cells, underscoring the requirement for regulated cortical tension in proper Cadherin-mediated adhesion^46^.

Of particular interest, NMII has been shown to play a key role in myoblast fusion in mouse, rat and drosophila by regulating cortical tension asymmetrically between the fusing cells^55,56^. In mouse myoblast fusion, for example, actors such as N-cadherin, ROCK, RhoA, Rac1, Cdc42, N-WASP and Arp2/3 have been shown to be involved ^47,57–59^. Similarly, ROCK inhibition has been reported to block membrane fusion in MDCK cells via an E-Cadherin–dependent mechanism, involving Arp2/3, Myosin II, RhoA, Cdc42 and Rac1^60,61^. These findings support our hypothesis that N-Cadherin, Arp2/3, Cdc42, RhoA, ROCK and NMII contribute altogether to TNT biogenesis by facilitating membrane fusion events.

By contrast, other studies have shown that NMII or ROCK inhibition enhances TNT-mediated transfer of α-synuclein in microglial cells^62^. While TNT biogenesis may vary between cell types, we propose that the formation of these structures involves conserved mechanisms, requiring precise modulation of key regulators due to their roles in pleiotropic cellular processes. Variations in protein expression patterns and activities between cells may further contribute to these differences. Notably, we also found that the stability of TNT-like connections correlates positively with N-Cadherin levels, echoing previous findings in HEK293 cells^34^.

NMII isoforms form complexes (NMII filaments) which bind actin filaments of different orientation and regulate their tension. They were thought to be none-processive before the recent observation of NMII-filaments processive motion in lamellipodia in cultured cells^50^. To our knowledge, NMIIA does not directly bind to cargoes such as intracellular vesicles or organelles known to transit through TNTs. Nevertheless, the observation of both processive motion and transfer of NMIIA filaments in our cells via the retrograde flow of actin suggests that TNTs could transfer bidirectionally (Fig. 6N-M). Indeed, other cargo-binding myosins such as Myosin XIX or Myosin Va could transfer between cells through processive motility or passively using the actin-treadmilling. Even non-motor scaffolding protein complexes tethering intracellular vesicles or organelles to actin could participate in TNT-mediated transfer. Due to the instability of TNTs, the success rate of either retrograde or anterograde transfer is expected to be inversely proportional to the length of the protrusion, and to be dependent on myosin processivity and speed. Supporting this assumption, NMIIA processive motion inside our TNT failed to travel the full protrusion length (∼2.7 μm out of ∼ 13 μm). Of particular interest, NMIIA processive motion appeared to be fast (∼ 290 nm/s) in TNTs (Fig. 6M-N) when compared to the average speed observed in lamellipodia in living cells (∼ 120 nm/s)^50^. Myosin speed can be differentially modulated by tropomyosin (Tpm) isoforms, filamentous proteins exhibiting different subcellular localization which can bind F-actin affecting its stability or affinity for actin-binding proteins^63^. The speed we measured appeared consistent with NMIIA speed observed *ex cellulo* on actin filaments decorated with Tpm 4.2 (∼160 nm/s without Tpm addition, ∼300nm/s on Tpm 4.2-decorated F-actin)^63,64^. This raises the question as to whether Tpm isoforms could regulate stability and material transfer within TNTs.

At the signaling level, we observed that Cdc42 expression increased in response to N-Cadherin overexpression and decreased following its knockdown. Cdc42 is a known mediator of Cadherin signaling^65^, and along with its downstream effectors IRSp53 and N-WASP, was essential for N-Cadherin–induced transfer in our system. Notably, this regulation occurred without apparent disruption to Cadherin homotypic binding, suggesting that these factors influence TNT-mediated transfer through mechanisms beyond adhesion stabilization. These findings are consistent with our previous work in mouse neurons showing that IRSp53 promotes functional TNT formation^41^ and with studies showing that Cdc42 and N-WASP are required for both TNT formation and vesicle transfer^41,51^. Taken together, our results support a model in which N-Cadherin overexpression enhances TNT-mediated transfer by upregulating Cdc42 expression, which, through its interaction with IRSp53 and N-WASP, promotes actin remodeling critical for TNT formation and function, acting in parallel to the increase in cell density and surface of cell-cell contact.

The precise mechanism by which these proteins promote transfer remains to be clarified. Cdc42, IRSp53, and N-WASP are well-established regulators of actin polymerization and filopodia formation^66,67,67^, suggesting a role in the elongation or stabilization of TNTs. However, these proteins are also implicated in various plasma membrane fusion processes^56,60,68,69^, a critical but understudied step in TNT biogenesis. These roles are not mutually exclusive: actin remodeling may drive protrusion formation, while localized ARP2/3 activation downstream of N-WASP could generate forces that support membrane fusion. While prior studies suggested that ARP2/3 inhibition enhances transfer by shifting the F-actin/branched-actin balance^41^, our findings support a model in which finely tuned, localized ARP2/3 activity downstream of Cdc42, IRSp53 and N-WASP is critical for fusion at membrane contact sites (Fig. 6M).

In summary, our data support an integrative model in which N-Cadherin promotes TNT-mediated contact-dependent transfer by coordinating membrane adhesion and cytoskeletal dynamics. This involves structural proteins (p120-Catenin, α-Catenin), regulators of cortical tension (ROCK, NMII), and actin modulators (Cdc42, IRSp53, N-WASP). These pathways act in concert with, but are mechanistically distinct from, density-induced increases in transfer. Given the multifunctionality of these proteins, future studies will require high-resolution tools to directly capture TNT formation and membrane fusion in real time. Approaches such as split-GFP or FRET-based reporters may help disentangle the dual roles of actin remodeling and membrane fusion in TNT biology.

## Material and Methods

### Cell lines, plasmids and transfection procedures

Human neuroblastoma (SH-SY5Y) cells were cultured at 37 °C in 5% CO2 in RPMI-1640 (Euroclone), plus 10% foetal bovine serum and 1% penicillin/streptomycin (gift from Simona Paladino, Department of Molecular Medicine and Medical Biotechnology, University of Naples Federico II, Naples, Italy). GFP N-Cadherin plasmid was available in the lab and was obtained from Sandrine Etienne-Manneville (Pasteur Institute, Paris, France) (53, 54), GFP NMIIA plasmids was a gift from Arnon Henn, Addgene (#11347). mEmerald α-Catenin plasmid was purchased from Addgene (#53982). To obtain clones that express GFP N-Cadherin or mEmerald α-Catenin, cells were transfected with the corresponding plasmid using Lipofectamine 2000 (Invitrogen) following the manufacture recommendations and selected with 300 ug/mL of geneticin for 10-14 days, changing the medium every 3-4 days. The pool of cells was seeded in 96-well plates through a limiting dilution in such a way that 0.5 cells are seeded per well, and after allowing them to grow, they were analysed and the clone overexpressing the protein of interest were selected. Human siRNA Oligo Duplex for N-Cadherin (SR300716) and α-Catenin (SR301060) and p120-Catenin siRNA (SR301064) were purchased from Origene. ON-TARGETplus Human BAIAP2 siRNA – SMARTpool (L-012206-02-0020), was purchased from DharmaconTM, Horizon Discovery. StealthTM RNAi Negative Contro Medium GC Duplex (452001) was purchased from Invitrogen. siRNA was transiently transfected to the cells through Lipofectamine RNAimax (Invitrogen) following the manufacture recommendations and the experiments are carried out in between 48 and 72 hours after the transfection.

### Sample preparation for visualization and quantification of the TNTs

*Wild-type* SH-SY5Y cells or overexpressing GFP or GFP N-Cadherin were trypsinized, counted, and 100.000 cells were plated overnight (O/N) on coverslips. Cells transfected with the corresponding siRNA were trypsinized and counted at 48 hours post-transfection and 100.000 cells were plated on coverslips (except for low density cells where only 60 000 cells were plated). 16 to 20 hours later (duration kept consistent within each experiment). Cells were fixed with specific fixatives to preserve TNTs first with fixative solution 1 (2% PFA, 0.05% glutaraldehyde and 0.2 M HEPES in PBS) for 15 min at 37 °C followed by a second fixation for 15 min with fixative solution 2 (4% PFA and 0.2 M HEPES in PBS) at 37 °C (for more information^39^). After fixation cells were washed with PBS and membrane was stained with conjugated Wheat Germ Agglutinin (WGA)-Alexa Fluor (1:300 in PBS) (Invitrogen) or Alexa Fluor-Phalloidin (1:300 in PBS) (Invitrogen) and DAPI (1:1000) (Invitrogen) for 15 minutes at room temperature, followed by 3 gentle washes with PBS. Finally, samples were mounted on glass slides with Aqua PolyMount (Polysciences, Inc.).

### Quantification of connected cells

Various Z-stacks images of different random points of the samples are acquired with either an inverted laser scanning confocal microscope LSM700 (Zeiss) controlled by the Zen software (Zeiss); or by a spinning disk Eclipse Ti2 inverted microscope (Nikon) equipped with a CSU- W1 spinning disk confocal scanning unit (Yokogawa), a 60× 1.4 NA oil immersion objective (Nikon Plan Apo VC), a fibre-coupled Coherent OBIS laser illumination system (405/488/561/640 nm), and a Prime 95B sCMOS camera (Teledyne Photometrics). Acquisition was controlled by Nikon’s NIS-Elements software. Z-stack slices of 0.5 μm thickness are encompassing the cells volume. Images are analysed following the morphological criteria of the TNTs: submicrometer wide membranous and/or actin-positive non-surface adherent structures connecting cells, first slices are excluded and only connections present in the middle and upper stacks are counted. Cells containing TNTs between them are marked as TNT-connected cells and by counting the number of cells that have TNTs between them and the total number of cells, the percentage of cells connected by TNTs is obtained. Analysis of the TNT-connected cells was performed in ICY software (https://icy.bioimageanalysis.org/) using the “Manual TNT annotation plugin”. At least 200 cells per condition were counted in each experiment. Images were processed with the ImageJ software.

### DiD transfer assay (co-culture assay)

DiD transfer assay is described elsewhere^39^. A co-culture is performed consisting of two populations of cells labeled differently: first, cells of interest (donors) are treated with Vybrant DiD (dialkylcarbocyanines), a lipophilic dye that stains the vesicles, 1:3000 (Thermo Fisher Scientific) in complete medium for 30 minutes at 37 °C (Life Technologies) before washing them with PBS. Then, these cells are co-cultured at a ratio of 1:1 with another population of cells (acceptors) marked in another colour (normally cells expressing soluble GFP or soluble mCherry) and grown for about 16 to 20 hours (duration kept consistent within each experiment), before fixing and staining them as described above for TNT-quantifications. For SH-SY5Y 50.000 donor cells are cocultured with 50.000 acceptor cells on glass coverslips within 24 well plates. The results are analyzed through microscopy described above and the final results are obtained by semi-quantitative analysis with the ICY or ImageJ software from calculating the percentage of acceptor cells with marked vesicles among the total number of acceptor cells. At least 100 acceptor cells per condition were counted in each experiment. Image montages were built afterward in ImageJ software. To measure the transfer in medium density and high density conditions. Secretion-mediated transfer was assessed in parallel to the DiD transfer assays, by two distinct methods.

Supernatant transfer protocol: DiD stained donors (50 000 cells) were grown in monoculture during 16 hours. Following that, supernatant was extracted to stimulate a monoculture of acceptors cells (50 000 cells) during 16 hours, before fixation and quantification of DiD positive acceptors.

ThinCerts protocol: DiD stained donors were grown (50 000 cells) on ThinCerts, pore size 1 μm (662610, Greiner Bio-one), placed on top of acceptors cells (50 000 cells) within the 24 well plate and cultured during 16 hour. ThinCerts’membranes does not allow cells to traverse, but enable small vesicles to pass freely.

### Chemical inhibitions

For both connection quantification and DiD transfer assay, drugs were added to all cells following seeding on the coverslips, before fixing them 16 hours later as described in the sample preparation for visualization and quantification of the TNTs and DiD transfer assay sections. ROCK inhibition was performed using Y-27632 (10 μM) (562822), purchased from BDBioscience; NMII inhibition using Bebbistatin, (10 μM) (B0560), purchased from MERCK; Cdc42 inhibition using ML141 (5 μM) (SML0407), purchased from MERCK; N-WASP inhibition using Wiskostatin, (10 μM) (681525), purchased from MERCK.

### Cell density measurement

Cell contours were traced in Fiji to measure the surface occupied by the cells. Cell nuclei were counted in parallel to obtain the cell density ratio for each biological replicate.

### Trypsin treatment experiment

Cell singularization by trypsinisation was adapted from ^42^. SH-SY5Y cells were plated the day before the experiment, seeding twice as many cells as under normal conditions, 800.000 cells per condition, since trypsin treatment would cause us to lose part of the cells that would detach in Ibidi μ-dishes (Biovalley, France) to favour cell adhesion with the substrate. 16 hours later the culture medium was replaced by 0.05% Trypsin/EDTA (Gibco), enough to cover the whole dish, for 90 seconds at room temperature. Immediately after these cells were fixed, stained, and analyzed exactly in the same way as described in “sample preparation for visualization and quantification of the TNTs”.

### Immunofluorescence

For immunofluorescence, 100.000 cells were seeded on glass coverslips and after O/N culture they were fixed with 4% paraformaldehyde (PFA) for 15 minutes at °C, quenched with 50 mM NH4Cl for 15 min, permeabilized with 0.1% Triton X-100 in PBS and blocked in 2% BSA in PBS. Primary antibodies used are: rabbit anti-N-Cadherin (ABCAM ref: ab76057), rabbit anti-N-Cadherin (Genetex ref: GTX127345) mouse anti-N-Cadherin (BD Biosciences ref: 610920), and rabbit anti-α-Catenin (Sigma ref: c2081) all of them at 1:1000 in 2% BSA in PBS during 1 hour. For p120-Catenin (Thermofisher 12180-1-AP) 1:100 in in 2% BSA in PBS overnight. After 3 washes of 10 minutes each with PBS, cells were incubated with each corresponding AlexaFluor-conjugated secondary antibody (Invitrogen) at 1:1000 in 2% BSA in PBS during 1 hour. For those experiments showing the actin cytoskeleton, cells were labeled with Rhodamine Phalloidin (Invitrogen) at 1:1000 in the same mix and conditions as the secondary antibodies. Then, cells were washed 3 times of 10 minutes each with PBS, stained with DAPI and mounted on glass slides with Aqua PolyMount (Polysciences, Inc.). Images were acquired with a confocal microscope LSM700 (Zeiss) and processed with the ImageJ software.

### Western blot

For Western blot cells were lysed with lysis buffer composed by 150 mM NaCl, 20 mM Tris, 5 mM EDTA, pH 8.0. Protein concentration was measured by a Bradford protein assay (Bio-Rad). Samples were boiled at 100 °C for 5 min and loaded in handcrafted 8% SDS-polyacrylamide gel or 4-12% Criterion™ XT Bis-Tris XT Precast Gels (Bio-Rad) and electrophoresed in 1X Tris/Glycine/SDS buffer (Bio-Rad) or 1X XT MOPS buffer (Bio-Rad) respectively for 1.5-2 hours at 90V. Proteins were transferred to 0.45 µm Nitrocellulose membranes (Bio-Rad) with 1X Tris/Glycine transfer buffer (Bio-Rad) for 1.5 hours at 90V in a cold chamber. Membranes were blocked in 5% non-fat milk in Tris-buffered saline with 0.1% Tween 20 (TBS-T) for 1 hour. Membranes were incubated O/N at 4 °C with the corresponding primary antibodies at 1:1000 in 5% non-fat milk TBS-T. Primary antibodies used for Western blot were: rabbit anti-N-Cadherin (ABCAM, ab76057), rabbit anti-N-Cadherin (Genetex, GTX127345) mouse anti-N-Cadherin (BD Biosciences, 610920), rabbit anti-α-Catenin (Sigma, c2081), mouse anti-IRSp53 (Invitrogen, MA537528) and mouse anti-α-tubulin (Sigma ref: T9026), rabbit anti-GAPDH (Sigma, G9545), rabbit anti-CDC42 (ABCAM, ab64533) rabbit anti-p120-Catenin (Thermofisher 12180-1-AP). Membranes were washed 3 times 10 minutes each with TBS-T and then incubated with the corresponding IgG secondary antibodies horseradish peroxidase-conjugated (GE Healthcare Life Sciences) between 1:1000 and 1:5000 for 1 hour at room temperature. Membranes were again washed 3 times 10 minutes each. Membrane protein bands were detected with Amersham™ ECL Prime Western Blotting Detection Reagent (Cytiva). Membranes were imaged using Amersham™ Imager 680 (GE Healthcare Life Sciences).

### Live Imaging

400.000 SH-SY5Y cells were plated the day before the experiment in Ibidi μ-dishes. After 16 hours of culture, live time series images were acquired with a 60 × 1.4NA CSU oil immersion objective lens on an inverted Elipse Ti microscope system (Nikon Instruments, Melville, NY, USA). Cells were labelled with 1:1000 dilution of conjugated WGA-Alexa Fluor in the corresponding media. Images were captured in immediate succession with one of two cameras, which enabled time intervals between 20 and 40 seconds per z-stack or between 50 and 70 seconds per z-stack when using two lasers. For live cell imaging, the 37 °C temperature was controlled with an Air Stream Stage Incubator, which also controlled humidity. Cells were incubated with 5% CO2 during image acquisition.

### Cell preparation for cryo-EM

Carbon-coated gold TEM grids (Quantifoil NH2A R2/2) were glow-discharged at 2 mA and 1.5–1.8 × 10-1 m bar for 1 minute in an ELMO (Cordouan) glow discharge system. Grids were sterilized under UV three times for 30 minutes at R. T. and then incubated at 37 °C in complete culture medium for 2 hours. 300,000 SH-SY5Y cells (RNAi N-Cadherin/α-Catenin, GFP N-Cadherin/mEmerald α-Catenin) were counted and seed on cryo-EM grids positioned in 35 mm Ibidi μ-Dish (Biovalley, France). After 24 hours of incubation, resulted in 3 to 4 cells per grid square. Prior to chemical and cryo-plunging freezing, cells were labeled with WGA (1:300 in PBS) for 5 min at 37 °C. For correlative light- and cryo-electron microscopy, cells were chemically fixed in 2% PFA + 0.05% GA in 0.2 M Hepes for 15 minutes followed by fixation in 4% PFA in 0.2 M Hepes for 15 minutes and kept hydrated in PBS-1X buffer prior to vitrification.

For cell vitrification, cells were blotted from the back side of the grid for 10 seconds and rapidly frozen in liquid ethane using a Leica EMGP system as we performed before (16).

### Cryo-electron tomography data acquisition and tomogram reconstruction

The cryo-EM data was collected from different grids at the Nanoimaging core facility of the Institut Pasteur using a Thermo Scientific (TF) 300kV Titan Krios G3 cryo-transmission electron microscopes (Cryo-TEM) equipped with a Gatan energy filter bioquantum/K3. Tomography software from Thermo Scientific was used to acquire the data. Tomograms were acquired using dose-symmetric tilt scheme, a +/-60 degree tilt range with a tilt step 2 was used to acquire the tilt series. Tilt images were acquired in counting mode with a calibrated physical pixel size of 3.2 Å and total dose over the full tilt series of 3.295 e-/Å2 and dose rate of 39,739 e-/px/s with an exposure time of 1s. The defocus applied was in a range of -3 to – 6 μm defocus.

Cryo-EM images c (Fig 1 B-D) were performed on a Tecnai 20 equipped with a field emission gun and operated at 200 kV (Thermo Fisher company). Images were recorded using SerialEM software on a Falcon II (FEI, Thermo Fisher) direct electron detector, with a 14 µm pixel size. The defocuses used were −6 µm.

The tomograms were reconstructed using IMOD (eTomo). Final alignments were done by using 10 nm fiducial gold particles coated with BSA (BSA Gold Tracer, EMS). Gold beads were manually selected and automatically tracked. The fiducial model was corrected in all cases where the automatic tracking failed. Tomograms were binned 2x corresponding to a pixel size of 0.676 nm for the Titan and SIRT-like filter option in eTomo was applied. For visualization purposes, the reconstructed volumes were processed by a Gaussian filter.

Quantitative manual measurements of iTNTs full extended, tip-closed, single TNTs were performed considering 10 cryo-EM slices and/or tomograms for control cells, 18 cryo-EM slices and/or tomograms for RNAi N-Cadherin, 45 for GFP N-Cadherin, 16 cryo-EM slices and/or tomograms for RNAi α-Catenin, 18 cryo-EM slices and/or tomograms for mEmerald α-Catenin, 10 cryo-EM slices and/or tomograms for co-culture of GFP N-Cadherin and RNAi α-Catenin.

### Cryo-EM N-Cadherin immuno-labeling

SH-SY5Y cells were plated on grids as described above. After incubation O/N at 37 °C, cells were fixed with PFA 4% for 20 min at 37 °C, quenched with 50 mM NH4Cl for 15 min, and blocked with PBS containing 2% BSA (w/v) for 30 min at 37 °C. Cells were labeled with a rabbit anti-N-Cadherin ABCAM 76057 antibody (1:200), followed by Protein A-gold conjugated to 10 nm colloidal gold particles (CMC, Utrecht, Netherlands). SH-SY5Y cells were then rapidly frozen in liquid ethane as above.

### RNA Extraction and cDNA Synthesis

Total RNA was extracted using the TRIzol reagent (Invitrogen #4374967), and RNA quality and quantity were assessed using the spectrophotometer DS-11 FX + DeNovix. 1 μg of total RNA was reverse transcribed using the High-Capacity cDNA Reverse Transcription Kit (Applied Biosystem #4368814) to generate complementary DNA (cDNA), following the manufacturer’s protocol.

### Quantitative PCR (qPCR)

qPCR was performed using the iTaq Universal SYBR Green Supermix (BIO-RAD #1725124) in a CFX384 Real-Time System C1000 Touch Thermal Cycler (BIO-RAD). Primers were designed to target the human BAIAP2 gene and a housekeeping gene (ACTB) as an internal control. Primer sequences were as follows:

- BAIAP2 forward: 5’AGCAGGAACTTTCGTGGTGA -3’
- BAIAP2 reverse: 5’-CTTCAGCGCAGCACTCAGAT -3’
- β-ACTIN forward: 5’-CCAACCGCGAGAAGATGA -3’
- β-ACTIN reverse: 5’-CCAGAGGCGTACAGGGATAG -3’

Each 10 µL reaction contained 5 µL SYBR Green master mix, 0.8 µM primers, and 2 µL cDNA (20 ng). Reactions were run in triplicate under the following cycling conditions: 95°C for 5 min, followed by 40 cycles of 95°C for 15 s, and 60°C for 30 s. Relative gene expression was calculated using the ΔΔCt method, normalizing the target gene to the housekeeping gene and comparing knockdown samples to the control. Results were plotted as fold change relative to the control. Statistical analysis was performed using GraphPad Prism, and significance was determined using unpaired t-test.

### Statistical analysis

Student’s t-test (for 2 groups) or One-Way ANOVA (for more than 2 groups) tests were applied. When significant differences were detected, Tukey’s Honestly Significant Difference (HSD) post hoc test was applied to identify specific group differences. A threshold of p < 0.05 was considered statistically significant. Normality and homogeneity of variances were assessed using the Shapiro–Wilk and Levene’s tests, respectively. All column graphs, Student’s t-test, and One-Way ANOVA statistical analysis were performed using GraphPad Prism version 9 software.

One-star significance was used to represent p-values below 0.05, two-star significance represents p-values below 0.01, three-star significance represents p-values below 0.001, four-star significance represents p-values below 0.0001.

## Acknowledgments

We thank all the laboratory members for useful discussion about theoretical and technical aspects, in particular Valeria Valente. We are grateful to R. Bouyssie, a member of the administrative staff of the Membrane Traffic and Pathogenesis department at Institut Pasteur. The NanoImaging Core at Institut Pasteur is acknowledged for support with image acquisition and analysis, particularly J.-M. Winter (NanoImaging Core at Institut Pasteur). We also acknowledge the Ultrastructural BioImaging platform of Institut Pasteur. We acknowledge Arnon Henn (Technion Israel Institute of Technology, Cellular Biology & Biophysics of Molecular Motors) for the plasmids GFP NMIIA.

## Author contributions

S.B., A.P., R.N.M., and C.Z. conceived the experiments. S.B., A.P., and R.N.M. prepared the figures and wrote the manuscript; S.B. and R.N.M. performed cocultures, TNT, and WB quantifications. A.P. performed cocultures, TNTs quantifications, and set up and performed all correlative, cryo-CLEM, and cryo-ET experiments, immunogold labelling by using TITAN cryo-EM, tomograms reconstruction, and quantitative analysis. A.S. performed EM acquisition by using Falcon F20. C. B. discussed experiments and generated the stable cell lines expressing aC-GFP and aC-RFP. S.B., A.P., R.N.M., C.B., and C.Z. discussed the results. All authors commented on the manuscript. C.Z. conceived the project, supervised all the work, and wrote the manuscript. C.Z. contributed to funding acquisition.

## Funding

This work was supported by the Fondation Recherche Médicale (FRM-EQU202103012692 and FRM MND202310017892), and Agence Nationale de la Recherche (ANR-20-CE13-0032 and ANR-23-CE16-0012-03) to C.Z. and The NanoImaging Core of Institut Pasteur was created with the help of a grant from the French government’s Investissements d’Avenir program (EQUIPEX CACSICE—Centre d’analyse de systèmes complexes dans les environnements complexes, ANR-11-EQPX-0008). We are grateful to late M. Michel, whose bequest to Institut Pasteur has made this project possible and Pasteur Joint International Research Unit «Neurodegenerative Diseases».

